# Neural feature spaces: characterizing the geometry of brain activity across development

**DOI:** 10.64898/2026.08.03.742613

**Authors:** Laurent Caplette, Rianne Haartsen, Saeideh Davoudi, Inga Sophia Knoth, Robert Leech, Emily Jones, Sarah Lippé

## Abstract

The human brain undergoes profound changes from childhood to adulthood. These changes are foundational to cognitive, affective and social development, and measuring them is essential for identifying atypical development. A common approach consists of analyzing metrics derived from brain activity, such as spectral power, aperiodic activity and signal complexity. However, individual metrics are sensitive to external factors unrelated to development, contributing to inconsistent findings in the literature. Moreover, these metrics are not independent; they show some degree of intrinsic redundancy, which itself evolves across development. Here, we propose a novel approach that focuses on the relationships between neural features rather than their values, defining a “neural feature space” that captures the geometry of brain activity. We analyzed the neural feature spaces of children (4−12 years) and adults (30−45 years), based on 128-channel EEG recordings during naturalistic movie viewing. Specifically, we computed a range of spectral, aperiodic and complexity features and quantified pairwise distances between them using latent variable modeling. Neural feature spaces were significantly different between age groups. Notably, high frequency bands were less differentiated in children and distances between spectral and complexity features differed. Crucially, distances were highly stable across tasks, in contrast to feature values, which varied substantially. These findings suggest that the geometry of neural feature spaces provides robust, interpretable markers of neurodevelopment, offering a complementary approach to feature-based analyses.

## 1 Introduction

The human brain undergoes profound structural and functional transformations across the lifespan. Among other changes, synaptic connections are gradually pruned (Huttenlocher, 1979; Sakai, 2020), axonal fibers are increasingly myelinated (Paus et al., 1999; Yakovlev & Lecours, 1967) and inhibitory interneuron networks are increasingly developed, leading to a greater excitation/inhibition ratio and more stable neural dynamics (Hensch, 2005; Uhlhaas et al., 2010). These developmental processes are reflected by changes in neural activity and can be assessed using electrophysiological methods such as electroencephalography (EEG), magnetoencephalography (MEG) and intracranial recordings. Investigating these changes is important, both to gain insights into the mechanisms of neurodevelopment and to identify deviations from a normal developmental trajectory.

A growing body of research has identified several neural features that are sensitive to development, including power in specific frequency bands, aperiodic activity parameters, and markers of signal complexity. Notably, the slope of 1/f-like aperiodic activity in the power spectrum flattens from childhood to adulthood (Cross et al., 2025; Merkin et al., 2023), high frequency power increases relatively to other frequencies (John et al., 1980; Miskovic et al., 2015), and brain activity becomes less predictable (increases in complexity; Davoudi et al., 2025; Lippé et al., 2009; McIntosh et al., 2010; Vakorin et al., 2011). Importantly, alterations of these features are also used as markers of abnormal development: for example, high frequency power is increased in Fragile X syndrome (Ethridge et al., 2017; Proteau-Lemieux et al., 2021), low frequency power is increased in Attention-Deficit/Hyperactivity Disorder (ADHD; Barry et al., 2003), and multiscale entropy, an indicator of signal complexity, is reduced in autism spectrum disorder (ASD; Bosl et al., 2011; Proteau-Lemieux et al., 2021).

Beyond differentiating various neurodevelopmental stages and disorders, such neural features are also sensitive to additional variables that may change across time (Davoudi et al., 2023). Feature values can vary depending on the task, the stimulus, and the participant’s emotional and cognitive state (e.g., Buszáki & Draguhn, 2004; Donoghue et al., 2020; Klimesch, 1997; McIntosh et al., 2010). This is useful in cognitive neuroscience studies to obtain information related to the transient brain state of a participant. However, when uncontrolled for in the experimental design, it can lead to contradictory results in the neurodevelopmental biomarkers literature (Donoghue, 2025; McVoy et al., 2019; Saby & Marshall, 2012). We posit that relationships between features are likely more stable within an individual than individual feature values. Any variation in those relationships may indeed be indicative of longer-term structural and functional transformations. For example, gamma power and neural complexity will both vary in response to stimuli viewed by a participant, but the relationship between gamma power and complexity is more likely to remain the same unless a more profound transformation in the organization of brain activity occurs.

In the present study, we propose to go beyond the values of individual neural features and analyze how a large number of features relate to each other across individuals. This set of inter-related features is defined as a *neural feature space*, which estimates the geometry of neural activity. The investigation of these relationships and how they change (or don’t) across development is relevant for several reasons. First, how various features are organized (including features from different tasks or preprocessing pipelines) could reveal how redundant they are, and therefore whether it is useful to investigate them (Dafflon et al., 2022). It could also lead to new insights into the neural mechanisms underlying those features. For example, some indicator of complexity being closely related to a particular frequency band would indicate that most of the variations in the predictability of the signal are driven by the presence of those oscillations. Relatedly, changes in those relationships would indicate a change in the mechanisms underlying a specific feature. As the brain evolves and cortical circuits mature, some neural features may reflect different processes. If a feature becomes driven by a different mechanism, we expect to see differences in how this feature relates to other features, irrespectively of whether its average value changes or not. Thus, changes in the organization of the neural feature space across development may provide new insights into neural development. Importantly, because the neural feature space is based on human-defined features, feature relationships remain easily interpretable, as opposed to features derived from black-box deep neural networks. Finally, the potential stability of feature distances across different tasks would represent a major asset and opportunity for multi-task and multi-site data aggregation.

In this study, we analyzed the neural feature spaces of neurotypical children (aged 4 to 12 years) and adults (aged 30 to 45 years) and how they differ. Specifically, the brain activity of 214 participants was recorded with a 128-channel EEG system while they were watching the animated movie *Rio* (2011). We then characterized i) the pairwise distances between spectral and complexity features; ii) how these distances changed across development; and iii) using a subset of those participants that had also performed an eyes-closed resting-state task, how stable they were across different tasks, compared to feature values.

## 2 Methods

### 2.1 Participants

In total, data from 270 participants was included in our study. All data was collected in the context of studies at the CHU Sainte-Justine Hospital (Montreal, QC, Canada). Within each dataset, all data was collected using the same EEG system and testing environment. All data collection was conducted according to procedures approved by the hospital’s Research Ethics Board, and informed consent was obtained by the participants or legal caregivers, in accordance with the Declaration of Helsinki.

Data from 214 participants was included in the main analyses (dataset 1a). There were 98 participants aged 4−12 years and 116 participants aged 30−45 years. Data was selected so that every participant had a minimum of 10 valid data segments (see below) and participants were excluded randomly so that both groups had an equal proportion of males and females (50%). None of the participants had a history of birth complication, health, neuropsychological or psychiatric conditions.

We then performed robustness analyses on two distinct datasets, each comprising a large range of ages (both children and adults). We first used data from 96 participants (overlapping with those included in our main analysis; dataset 1b) that had data from two different tasks (39 females; mean age = 9.75, S.D. = 4.42). We also ran the same analysis on a separate dataset (dataset 2) that was recorded with a different EEG system in a different environment and with partly different tasks. After the exclusion of participants with less than 10 valid data segments, data from 56 participants (32 females; mean age = 32.96, S.D. = 16.94) remained.

### 2.2 Design considerations and sample size

Because of our novel analytical technique, expected effect sizes were unknown. Since our study was a reanalysis of data collected in the context of other studies, we chose to maximize our statistical power by using all data available that conformed to our inclusion criteria (see Participants).

### 2.3 EEG recording and procedure

For the main analyses (datasets 1a and 1b), EEG data was acquired using a 128-channel EGI HydroCel Geodesic Sensory Net and a Net Amps 300 EEG amplifier (MagStim EGI, Eugene, OR, USA) at a 1000 Hz sampling rate. A G4 Macintosh computer running NetStation EEG software (Version 4.5.4) was used to synchronize recordings with experimental events and to store the data. Resting-state data was acquired for around 5 minutes on average and up to 15 minutes. A clip from the animated movie *Rio* (2011) was shown full-screen during recording. The movie was projected on a Tobii T120 Eye Tracking screen with a resolution of 1024 × 1280 pixels and a refresh rate of 60 Hz. Impedances were kept under 40 kΩ and the vertex was used as online reference during recording. For additional analyses, we used data from participants that had performed both the above task and an additional resting-state task in which they had their eyes closed (with the same experimental setup; dataset 1b).

We performed supplementary analyses on a separate dataset (dataset 2), acquired using a 128-channel EGI HydroCel Geodesic Sensory Net and a Net Amps 400 EEG amplifier (MagStim EGI, Eugene, OR, USA) at a 1000 Hz sampling rate. An iMac computer running NetStation EEG software (Version 5) was used to synchronize recordings with experimental events and to store the data. A first set of resting-state data was acquired for around 5 minutes while a clip from the movie *Rio* was shown (RIO task), and then for another 5 minutes while 4 abstract screensaver-like videos of moving shapes (lasting 1 minute each, with intermissions of around 20s in between) were shown (SHAPES task). *Rio* was shown full-screen and abstract videos were relatively small, covering approximately 9.1 x 6.2 degrees of visual angle. The stimuli were projected on an ASUS VG248QE screen with a resolution of 1920 × 1080 pixels and a refresh rate of 144 Hz. Impedances were kept under 40 kΩ and the Cz electrode was used as online reference during recording.

### 2.4 EEG preprocessing

Preprocessing was performed using the HAPPE (version 3) pipeline (Gabard-Durnam et al., 2018) in Matlab R2023a for all data. This pipeline was developed specifically to preprocess infant and child data (but here was used for all data including from adult participants). Specifically, (1) data was filtered between 1 and 80 Hz using EEGlab’s Finite Impulse Response filter; (2) 20 frontal and occipital channels at the outer edge of the net were excluded due to them being more prone to noise from head movements; (3) electrical noise around 60 Hz (± 3 Hz) was removed using the multi-taper regression method implemented by the CleanLine approach in EEGlab (Mullen, 2012); (4) bad channels with high impedances or displacement during recording were removed and data was interpolated from nearby channels; (5) a wavelet-enhanced ICA was performed to remove major artifacts and improve the subsequent ICA decomposition of the data; (6) ICA with automatic component rejection was performed to remove remaining artifacts; (7) data was segmented into 5-second epochs, with epochs in which at least one channel exceeded an amplitude of -150 μV or 150 μV were removed; (8) data was re-referenced to an average reference.

### 2.5 EEG feature extraction

From the preprocessed EEG data, we computed various features that reflect important aspects of brain activity that have been related to neural development and various neurodevelopmental disorders (Bosl et al., 2011; Cross et al., 2025; Davoudi et al., 2025; Lippé et al., 2009; Miskovic et al., 2015; Proteau-Lemieux et al., 2021). Feature extraction was performed in Python, for each electrode and half of the data within each participant; participants for which the extraction of any feature failed were excluded (3 participants). SpecParam (formerly known as FOOOF; Donoghue et al., 2020) was used to estimate aperiodic activity parameters and power. In this approach, the power spectrum is conceived as a combination of an aperiodic 1/f-like component and a variable number of periodic oscillations. Specifically, power spectral density from 1 to 80 Hz (at a resolution of ∼0.24 Hz) was first estimated for each segment using Welch’s method. Then, power spectra were averaged within each data half. Finally, a FOOOF model (default parameters) was fitted to each averaged spectrum to estimate the aperiodic component parameters, i.e., an offset representing the height of the aperiodic spectrum and an exponent representing its slope. Power adjusted for aperiodic activity (henceforth, periodic power) was then estimated for the delta (1−4 Hz), theta (4−8 Hz), alpha (8−13 Hz), beta (13−30 Hz), low gamma (30-57 Hz) and high gamma (63−80 Hz) frequency bands by averaging spectral density within these bands after having removed the aperiodic component.

Various complexity features were estimated for each data segment, and values averaged across segments for each data half. MNE-Features was used to estimate fractal dimension, Hurst exponent and sample entropy, while NeuroKit2 was used to estimate multiscale entropy (MSE). Fractal dimension quantifies the self-similarity and space-filling characteristics of a curve (here, time series) and its value varies between 1 and 2. For example, a flat line has a fractal dimension close to 1 while a curve with lots of repeating patterns at multiple scales fills more space and has a fractal dimension closer to 2. Fractal dimension was estimated with Higuchi’s method, which is reliable and appropriate for nonstationary data (Shi, 2018; Lau et al., 2022). Higuchi’s algorithm reconstructs many time series by downsampling the original signal every *k* samples, for different *k* values, estimates the length of each resulting curve, and estimates the fractal dimension from there. The Hurst exponent quantifies long-range temporal autocorrelation. Its value ranges between 0 and 1, although it usually varies between 0.5 and 1 for EEG signals. A Hurst exponent of 0.5 indicates an absence of temporal autocorrelation, whereas a higher value indicates a positive autocorrelation, with past time points being positively correlated to future time points. Sample entropy is an information-theoretic measure of the complexity of the signal, with higher values representing a more complex, less predictable signal. MSE is an extension of sample entropy evaluating the predictability of the signal at multiple scales. The time series is downsampled at multiple scales and sample entropy is computed for each scale. Low-scale and high-scale complexity indices (CI) were then computed by integrating sample entropy across low (1^st^−20^th^) and high (21^st^−40^th^) scales separately.

Finally, all feature values were averaged into 8 ROIs of electrodes (occipital, frontal, central, temporal left, temporal right, parietal left, parietal right, whole-brain). Henceforth, a “feature” refers to a specific value for a given ROI and feature type (e.g., Hurst exponent for the frontal ROI). A total of 104 features (8 ROIs × 13 feature types) per participant, with 2 values each (for each data half), were thus entered into subsequent analyses.

### 2.6 Neural feature spaces

We computed pairwise distances between all features across participants of each age group. To ensure that distances were independent of average feature magnitude and sign, our distance of choice was the absolute Pearson correlation distance, defined as:

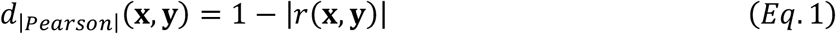

where *r* is the Pearson correlation coefficient. However, comparing observed correlation coefficients to each other can be problematic (Diedrichsen et al., 2018; Guggenmos et al., 2018; Walther et al., 2016). Indeed, correlation coefficients are biased estimators that are attenuated by noise and noise level might vary between groups and across features. Note that, being aggregate measures computed from reasonable amounts of data, our feature values were on average highly reliable, and similarly reliable between groups (mean *r* between data halves was 0.90 for both groups). Nevertheless, to ensure accurate inferences, we used latent variable modeling to approximate the disattenuated correlations between latent features and statistically compare distances derived from them (Archer et al., 2008; Hancock, 1997; Wang, 2010).

Specifically, for each pair of features, we fit a simple multi-group latent variable model to the data using the lavaan toolbox in R (Rosseel, 2012). For each group, we modelled two latent variables (the two features under investigation) and two indicators per variable (the two values from distinct data halves). Each latent variable was related to its indicators with equal loadings; this loading was a parameter estimated in the model fitting. The variance of each latent variable, as well as the covariance between both latent variables, were also estimated parameters. For each latent construct, the residual variances of each indicator were also estimated parameters with equal value. All parameters were allowed to be different across groups. From the latent covariances and variances, we computed the correlation for each group and clamped it between −1 and 1 (in case small variances led to a misestimation of the coefficient beyond normal boundaries). We also defined additional parameters derived from those correlations for statistical testing (see below). The models were fit using robust maximum likelihood and Huber-White robust standard errors were computed using the delta method. Given the very different scales of different features, data were standardized (across groups) prior to fitting to favor model convergence. All models converged (median CFI = 0.96, median SRMR = 0.04). For some models, standard errors (and therefore p-values) of some parameters could not be computed (Children group correlations: 0.11% of models; Adult group correlations: 0.39% of models; Difference of correlations: only 1 model, 0.04%); in those cases, results were deemed nonsignificant with a p-value of 1.

Because the absolute value function is not differentiable, we could not directly use it to define a parameter in our model and obtain a p-value for it using the delta method. However, we could perform equivalent tests by defining additional parameters. Specifically, we defined a “sub” parameter for each group as 1 minus the latent correlation for that group, an “add” parameter for each group as 1 plus the latent correlation for that group, a “diff” parameter as the difference between the correlations of both groups and a “sum” parameter as the sum between them. To assess whether distances in an individual group were significantly different from 0, we took the maximum p-value of the “sub” and “add” parameters for that group. This test is equivalent to testing 1 – abs(r) and at least as conservative: indeed, if both parameters are different from 0, it entails that the correlation is different from both 1 and –1, and hence the distance is guaranteed to be greater than 0. To test whether distances in an individual group were significantly different from 1, we used the two-tailed p-values associated to the test of the correlations against 0. This test is equivalent to a one-tailed p-value for the absolute value of the correlation against 0 (and hence to the one-tailed p-value for our distance against 1), given that the standard two-tailed test for a correlation depends only on the correlation’s magnitude. To test whether distances were different between groups, we took the maximum of the “diff” and “sum” p-values. This test is conservative and equivalent to testing the difference between the absolute values of the correlations: indeed, if correlations were of equal magnitude but opposite sign, they would sum to 0 (and hence the “sum” parameter would not be significant, even if the “diff” is). Finally, p-values were corrected for multiple comparisons using the Benjamini-Hochberg procedure to control the False Discovery Rate.

To visualize the neural feature space of each group, we used the estimated correlations and computed the absolute Pearson correlation distances from them. We then reduced the high-dimensional matrix of all pairwise distances to a set of two-dimensional coordinates using metric multidimensional scaling (MDS). This technique enables finding coordinates in a given number of dimensions for each element such that distances in that lower-dimensional space stay as close as possible to the original distances. Note that distances can normally only be interpreted within each space and not across spaces. Since MDS coordinates are arbitrary (only the relative positions are meaningful), we aligned the spaces of both groups to each other as well as possible without distorting them using Procrustes transformations (translation, scaling, reflection and rotation).

To visualize the uncertainty around MDS coordinates, we recomputed distances under a bootstrap resampling scheme. Due to computational constraints, we could not run an additional model fit each time but instead used predicted latent scores for each feature from the already fit models. For each feature, scores were averaged across all models in which that feature was included. Correlations between latent scores are roughly equivalent to latent correlations as estimated by a model. Feature distances and MDS coordinates estimated this way were highly similar to the feature distances and MDS coordinates (after Procrustes transformation) estimated by the models (all *r* > .99). We then resampled participants with replacement 10,000 times within each group and recomputed distances from their latent scores. We performed MDS, aligned the resulting space to the observed space of the respective group using a Procrustes transformation, and recorded the resulting coordinates. The bootstrap standard errors of the x and y coordinates are illustrated.

### 2.7 Robustness analyses

We compared the across-tasks robustness, or stability, of the feature distances (computed across participants) to that of the feature values (averaged across participants). We first compared data from the *Rio* task to data from an eyes-closed task, on neurotypical children and adults. As a supplementary analysis on additional data (see *Participants*), we compared the *Rio* task to an abstract videos task. Since distances are bounded estimates, their magnitude cannot easily be compared to feature values; we therefore compared only the relative values by computing a correlation of feature values across tasks and a correlation of feature distances across tasks. However, since feature values are all on different scales, a correlation computed naively would be primarily driven by those differences in scales. Therefore, we normalized those values using separate participants prior to computing distances and correlations. Moreover, due to computational constraints (especially given statistical testing), we fit the model only once per pair of features and obtained latent correlations by computing correlations on predicted latent scores, which results in highly similar estimates (*r* > .99, see above).

Specifically: (1) we fit a model and predicted latent scores for each pair of features and task separately (with the same free parameters as our main model except there is only one group); (2) for each task and feature, latent scores were averaged across all models in which that feature was included; (3) we randomly divided participants into equal training and test sets; (4) within the training set, we averaged both observed data and latent scores across tasks (observed data was also averaged across halves) and computed the mean and standard deviation across participants, to get an independent estimate of the scale of each feature (for observed data and latent scores separately); (5) data and latent scores from the test set were centered and scaled using the training set means and standard deviations; (6) feature distances (i.e., absolute Pearson correlation distances) for each task were estimated using the standardized latent scores; (7) feature values for each task were estimated by averaging standardized observed feature values across participants; (8) an estimate of the distance stability was obtained by computing the across-tasks correlation of the feature distances; (9) an estimate of the value stability was obtained by computing the across-tasks correlation of the feature values; (10) the stability measures were transformed to Fisher-Z values and their difference was computed; (11) this analysis (steps 6-10) was repeated while resampling participants within the test set with replacement 10,000 times to establish a bootstrap distribution of the difference in stability; (12) this whole procedure (steps 3-11) was repeated with 1,000 random splits of participants; (13) all estimates were averaged across the 1,000 random splits; (14) the observed stability difference and its bootstrap distribution were standardized using the standard deviation of the bootstrap distribution; (15) the presence of 0 within the 95 and 99.9 central percentiles of the bootstrap distribution was assessed to determine statistical significance (at α=0.05 and α=0.001).

## 3 Results

### 3.1 Neural feature spaces

The high-dimensional neural feature space of each group can be reduced to two dimensions with minimal distortion and subsequently visualized (Figure 1). When analyzing each neural feature space, we first observe that for most features, the different ROIs largely cluster together, which is expected given the high spatial correlation present in EEG (although some ROIs occasionally steer away from their main cluster). This suggests that sampling multiple ROIs is likely unnecessary in most cases. For both groups, theta, alpha and beta power are relatively close to each other, while low and high gamma form a distinct cluster. The Hurst exponent is close to the high-scale complexity index, while sample entropy is close to the low-scale complexity index and the aperiodic exponent; fractal dimension is in between. Overall, almost all distances were significantly different from 0 (99.49% of distances in children and 97.96% of distances in adults; *p* < .05, one-tailed, FDR-corrected), which is expected, given that distinct features rarely covary in exactly the same way, with perfect redundancy. This entails that most features do not encode the exact same information. Of the few features that had a near-0 distance, most were of the same feature type for different brain regions (e.g., temporal and parietal theta power). Additionally, we tested whether distances were significantly different from 1 (i.e., whether correlations between features were non-zero). This was the case for 64.88% of distances in children and 65.78% of distances in adults (*p* < .05, one-tailed, FDR-corrected), indicating that most features were not perfectly independent from each other. Overall, we observe that spectral features cover a large of the space, but that complexity features bring additional information.

**Figure 1.**
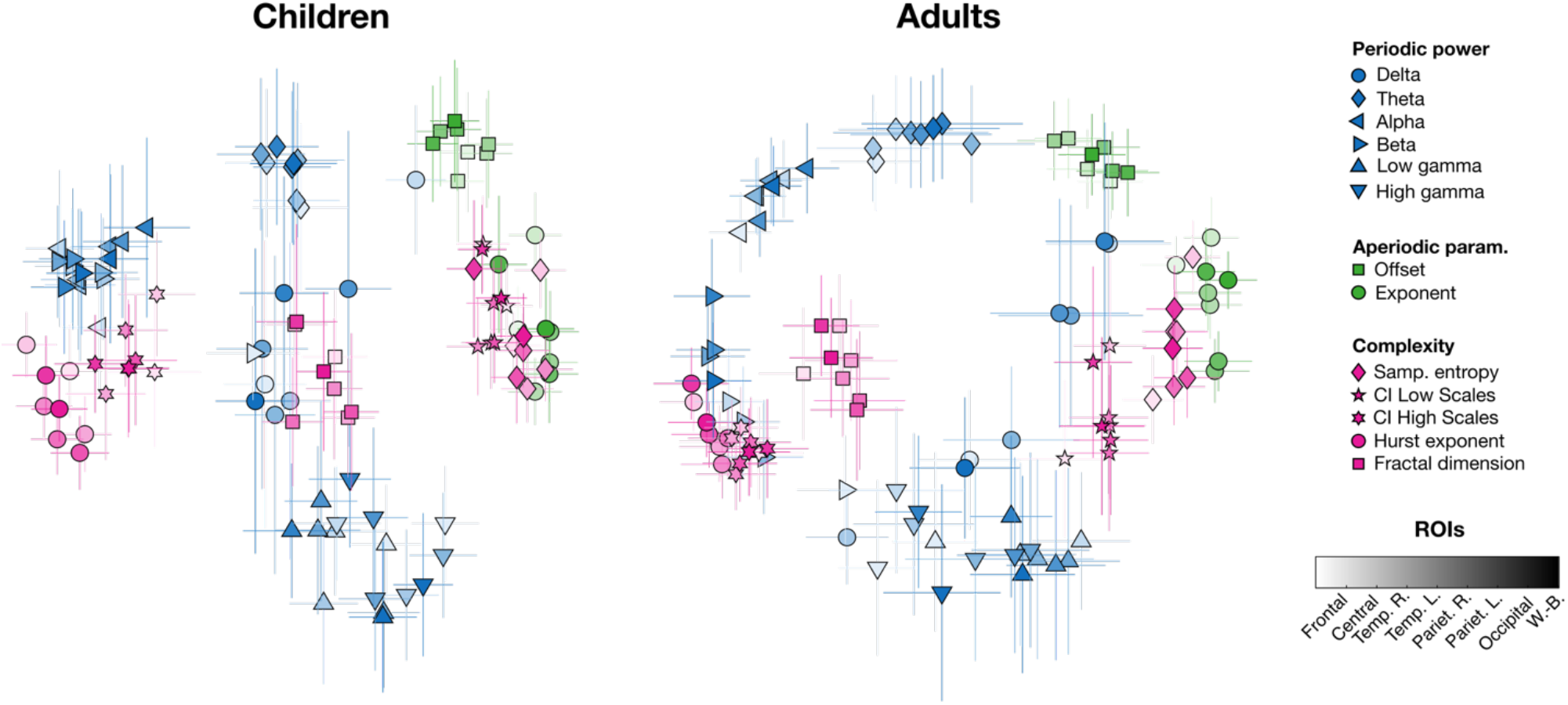
Neural feature spaces of children (4−12 years) and adults (18−35 years), reduced to two dimensions. Only relative distances can be compared between groups in this visualization; axes are not meaningful. Each symbol/color combination corresponds to a different feature type, while color saturation indicates ROI (legend on the right). Error bars indicate bootstrap standard errors in the 2D coordinates.

Although there were similarities between the spaces of both groups, we also found many differences. Overall, 107 distances were significantly different between the groups (*p* < .05, two-tailed, FDR-corrected; Figure 2). Sixty-two of those were greater in children than adults. Of note, the Hurst exponent was generally more distant to complexity indices, delta power (in the temporal ROI) and beta power in children. Moreover, delta power in children was more distant to both high gamma and the low-scale CI; and the high-scale CI was more distant to high gamma, fractal dimension, sample entropy and the low-scale CI in children. On the flip side, 45 distances were smaller in children than adults. Low and high gamma were closer to the low-scale CI; fractal dimension was closer to the aperiodic exponent and the low-scale CI; frontal alpha was closer to high-scale complexity; and low and high gamma were closer to each other. Overall, these results indicate that feature distances can discriminate age groups. Moreover, the specific differences that were found indicate that neural oscillations and complexity are differentially related across development.

**Figure 2.**
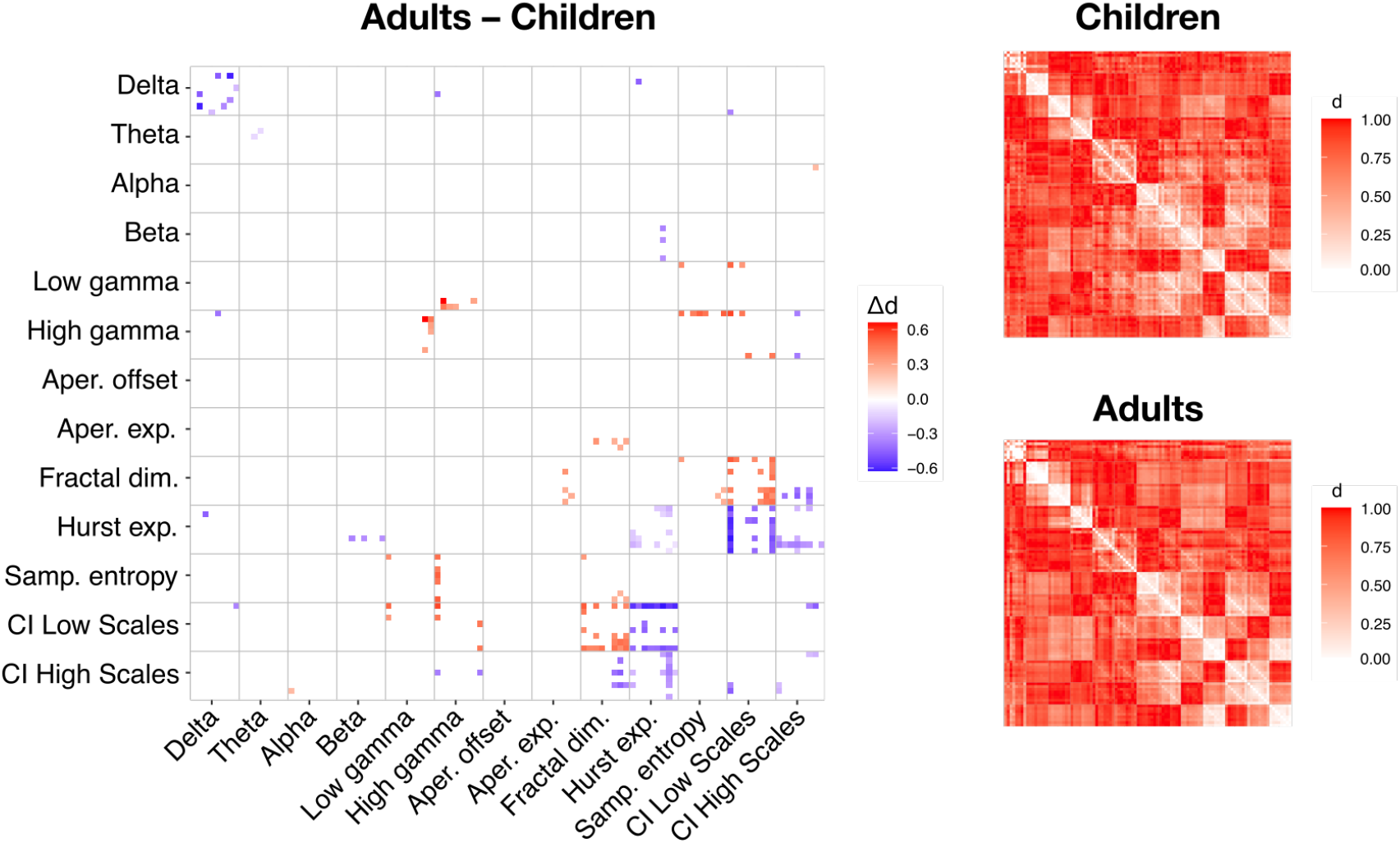
Illustration of significant differences of distances between children and adults (*p* < .05, two-tailed, FDR-corrected); distances that are greater in adults are visible in red, while distances that are greater in children are visible in blue. Smaller matrices on the right show all distances in each group (with features in the same order). Each element of each matrix refers to the specific distance between corresponding features on the *x* and *y* axes (matrices are symmetric along the diagonal). Each pair of features is represented as a small 8 x 8 square within each matrix. Within these squares, each pair of ROIs is represented as a distinct element, from top to bottom and from left to right: frontal, central, right temporal, left temporal, right parietal, left parietal, occipital and whole-brain.

### 3.2 Robustness to task and stimulus

We verified whether distances between features were more robust to the task and stimulus than the feature values were. To do so, we analyzed the data from 96 participants (dataset 1b; both children and adults; mean age of 9.75 years) that had performed both the task described above (RIO task) and a resting-state task with eyes closed (REST task). We correlated normalized feature values and distances across tasks (see Methods). We observed that feature distances were much more stable across tasks than feature values. In fact, feature values were anti-correlated (which is to be expected, especially e.g. for alpha power, given the very different sensory inputs in both tasks – eyes open vs eyes closed), while feature distances were highly positively correlated (*r* = −0.63 vs *r* = 0.80; *Z* = 205.85; *p* < .001, two-tailed; Figure 3a-b).

**Figure 3.**
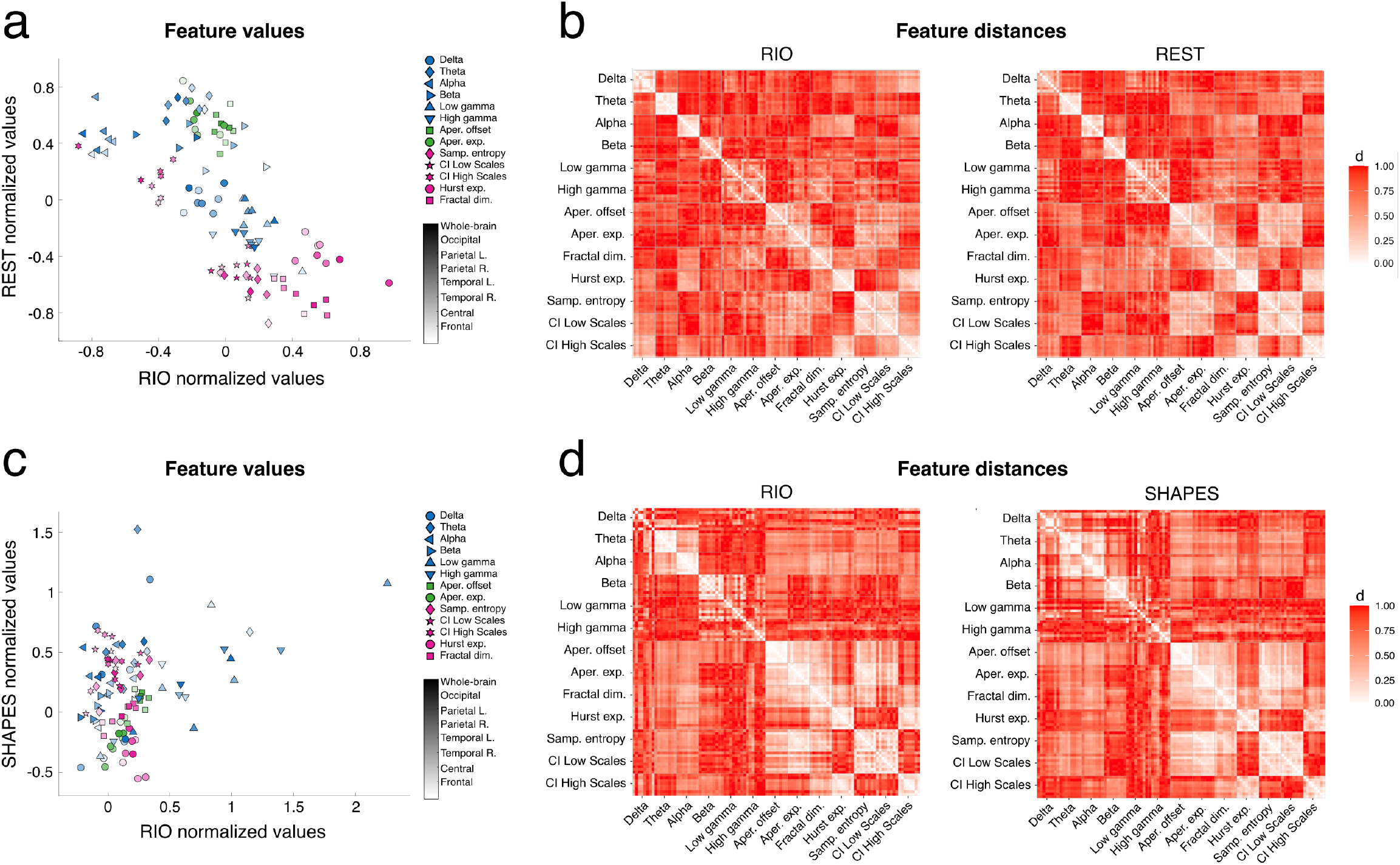
Feature values and distances across tasks. **A)** Normalized feature values for the RIO (*x* axis) and REST (*y* axis) tasks in dataset 1b. Legend of feature types and ROIs is on the right. **B)** Feature distances for the RIO (left panel) and REST (right panel) tasks in dataset 1b. Matrices are symmetric along their diagonals. **C)** Same as **A** for the RIO and SHAPES tasks in dataset 2. **D)** Same as **B** for the RIO and SHAPES tasks in dataset 2. Note that, given that multiple normalizations were performed using random subsets of participants, we displayed here the ones that resulted in the correlation values closest to the reported mean of correlation values.

We replicated this result on data from 56 additional participants (dataset 2; also both children and adults; higher mean age of 32.96 years) that was recorded on a different EEG system. For this dataset, the two tasks to be compared were more similar, in the sense that participants had their eyes open during both: in addition to a clip from *Rio*, participants were shown screensaver-like videos of moving abstract shapes (SHAPES task). When comparing the stability of feature distance estimates and feature value estimates across these tasks, we found that feature values were moderately positively correlated, while feature distances were highly positively correlated (*r* = 0.33 vs *r* = 0.82; *Z* = 84.16; *p* < .001, two-tailed; Figure 3c-d).

## 4 Discussion

We developed a novel approach to characterize differences in the brain activity of children and adults. Instead of focusing on specific biomarkers, we characterized exhaustively the space of neural features in each group. Rather than focusing on how the values of features differ between the groups, we used latent variable modeling and analyzed how the relationships between the features differed.

Across both groups, distances between features revealed a coherent organization that aligns with plausible temporal and spectral properties of EEG signals. The Hurst exponent and high-scale CI showed a strong association, consistent with both indexing long-range temporal dependencies and thus capturing aspects of slow dynamics. In contrast, sample entropy was highly correlated with low-scale complexity, suggesting that it primarily reflects irregularity at shorter timescales, likely driven by higher-frequency components of the signal. Notably, sample entropy also exhibited a close relationship with the aperiodic slope, indicating that signal predictability is tightly linked to the properties of aperiodic activity. In terms of distances with spectral features, high-scale complexity was more closely aligned with lower-frequency oscillations, particularly in the theta and beta ranges, which is expected given that high-scale complexity mostly reflects the predictability of the signal at longer time scales. Moreover, the aperiodic offset was closer to the delta and theta frequency bands, which is consistent with the overall level of the spectrum being more influenced by low frequencies. Importantly, all spectral features, including aperiodic parameters and periodic oscillations, remained largely independent from each other, reinforcing the view that periodic and aperiodic components capture complementary, non-overlapping aspects of brain activity. Overall, almost all features were significantly distant from each other, indicating that, despite their obvious relationships, no two features were completely redundant, and suggesting that different features reflect partially distinct underlying mechanisms. These results highlight the importance of considering the joint structure of features rather than solely interpreting them in isolation.

When looking at differences between groups, we can observe a marked reorganization in the relationships between spectral and complexity features from childhood to adulthood. In children, frontal low gamma power is closely related to short timescale complexity, but this is much less the case in adults. This is consistent with the notion that gamma activity is less stable and more transient early in development, likely reflecting immature excitation-inhibition balance, particularly in frontal regions, whereas in adulthood gamma becomes more tightly regulated and functionally specific. More broadly, the association between gamma power and complexity indices was reduced in adults compared to children, suggesting that high-frequency activity becomes less directly tied to local signal irregularity with maturation. The relationship between the Hurst exponent and the multiscale entropy CIs was also different across development. While both features are usually negatively correlated in EEG data, due to the way they are defined (a more predictable signal will have higher Hurst exponent but lower CI), this correlation is less strong in children (i.e. the distance is greater). This potentially reflects the contribution of a broader range of frequencies into long-range complexity in adults. In addition, developmental changes were evident within the gamma band itself: in children, low gamma and high gamma power were closely related, while they were more distant in adulthood. This suggests that gamma activity is more broadband and less differentiated early in development. These results are consistent with prior work showing that stimulus-induced gamma is more broadband in children (Rhodes et al., 2025). With maturation, these bands become more distinct, likely due to the progressive refinement of inhibitory interneuron networks. Computational modeling further supports this interpretation, indicating that a developmental shift toward stronger inhibition relative to excitation in superficial cortical layers contributes to more band-limited gamma activity in adulthood, independently of changes in aperiodic slope (Rhodes et al., 2025).

Analyses of stability across tasks indicate that feature distances constitute markers of neurodevelopment more robust to the task compared to raw feature values. While individual feature estimates showed substantial variability across tasks, the relative structure of features, captured in our neural feature spaces, remained comparatively stable. This suggests that task- and stimulus-dependent fluctuations primarily affect the absolute magnitude of features, whereas their organization is preserved, thus facilitating cross-study comparability. Moreover, the robustness of feature distances raises the possibility of combining data from distinct tasks or conditions, thereby increasing statistical power and generalizability. Finally, focusing on distances rather than raw values may offer deeper insights into brain mechanisms that remain stable across tasks and robust to short-term external influences. This increased stability is not guaranteed and will depend on the specific features considered, but in general more abstract constructs are likely to be more stable than the simpler parts they are abstracted from.

Our study is limited by the specific age groups and the relatively modest samples (214, 96 and 56 participants) that were available to us. In future studies, we hope to expand this approach and assess more continuous changes across development, from early childhood to late adulthood. Our approach can also naturally be extended to the comparison of various neurodevelopmental disorders and genetic conditions. It would be particularly interesting to assess whether multiscale entropy and gamma power are differentially related in ASD. Moreover, as mentioned earlier, statistical power could be increased by combining multiple tasks in the same neural feature spaces, which seem to be minimally affected by task and stimulus. Cognitive features, such as questionnaire items and performance indicators in various tasks, could also be included in these spaces, in order to provide a more comprehensive phenotyping. Finally, we note that some of our results may be affected by activity from non-neural sources (e.g., movement) differing between groups. However, several layers of preprocessing were used to prevent this as much as possible (bad channel rejection, wavelet-enhanced ICA, regular ICA, amplitude-based segment rejection). In the future, we hope that our results can be replicated with intracranial recordings, which are less affected by external artifacts, to provide additional validation.

In summary, our approach brings to the forefront new dimensions to investigate in the characterization of neurodevelopment. In addition to investigating common neural features, we favor the adoption of a holistic approach that characterizes the overall geometry of these features in a neural feature space. These neural feature spaces bring new information allowing us to discriminate between different neurodevelopmental stages and they are robust to the task and the stimulus, paving the way for new promising biomarkers.

## Acknowledgements

We wish to thank all participants and families who took part in this study.

## Funding

This work was supported by a grant from the Simons Foundation (#986609, SFARI GAIINS; E.J. and S.L.), a grant from NSERC (#RGPIN-2023-04848; S.L.) and the Canada Research Chair on Pediatric Neurodiversity (CRC-2024-00030; S.L.).

## Code availability statement

All original code used for data analysis will be shared on Github upon publication.

## Author contributions (CRediT)

Laurent Caplette: Conceptualization, Data curation, Formal analysis, Methodology, Software, Writing – original draft, Writing – review and editing

Rianne Haartsen: Methodology, Software, Writing – review and editing

Saeideh Davoudi: Data curation, Methodology, Software, Writing – review and editing

Inga Sophia Knoth: Investigation, Project administration, Writing – review and editing

Robert Leech: Writing – review and editing

Emily Jones: Methodology, Validation, Funding acquisition, Writing – review and editing

Sarah Lippé: Supervision, Validation, Funding acquisition, Writing – review and editing

## Notes

### Competing Interest Statement

The authors have declared no competing interest.

## References

Archer, K. J., Dumur, C. I., Taylor, G. S., Chaplin, M. D., Guiseppi-Elie, A., Buck, G. A., … & Garrett, C. T. (2008). A disattenuated correlation estimate when variables are measured with error: Illustration estimating cross-platform correlations. Statistics in Medicine, 27(7), 1026–1039.

Barry, R. J., Clarke, A. R., & Johnstone, S. J. (2003). A review of electrophysiology in attention-deficit/hyperactivity disorder: I. Qualitative and quantitative electroencephalography. Clinical neurophysiology, 114(2), 171–183.

Bosl, W., Tierney, A., Tager-Flusberg, H., & Nelson, C. (2011). EEG complexity as a biomarker for autism spectrum disorder risk. BMC Medicine, 9(1), 18.

Buzsáki, G., & Draguhn, A. (2004). Neuronal oscillations in cortical networks. Science, 304(5679), 1926–1929.

Cross, Z. R., Gray, S. M., Dede, A. J., Rivera, Y. M., Yin, Q., Vahidi, P., … & Johnson, E. L. (2025). The development of aperiodic neural activity in the human brain. Nature human behaviour, 9, 2548–2563.

Dafflon, J., F. Da Costa, P. Váša F., Monti, R. P., Bzdok, D., Hellyer, P. J., … & Leech, R. (2022). A guided multiverse study of neuroimaging analyses. Nature Communications, 13(1), 3758.

Davoudi, S., Arango, G. L., Deguire, F., Knoth, I. S., Thébault-Dagher, F., Reh, R., … & Lippé, S. (2025). Electroencephalography estimates brain age in infants with high precision: Leveraging advanced machine learning in healthcare. NeuroImage, 312, 121200.

Davoudi, S., Schwartz, T., Labbe, A., Trainor, L., & Lippé, S. (2023). Inter-individual variability during neurodevelopment: an investigation of linear and nonlinear resting-state EEG features in an age-homogenous group of infants. Cerebral Cortex, 33(13), 8734–8747.

Diedrichsen, J., Yokoi, A., & Arbuckle, S. A. (2018). Pattern component modeling: A flexible approach for understanding the representational structure of brain activity patterns. NeuroImage, 180, 119–133.

Donoghue, T. (2025). A systematic review of aperiodic neural activity in clinical investigations. European Journal of Neuroscience, 62(7), e70255.

Donoghue, T., Haller, M., Peterson, E. J., Varma, P., Sebastian, P., Gao, R., … & Voytek, B. (2020). Parameterizing neural power spectra into periodic and aperiodic components. Nature neuroscience, 23(12), 1655–1665.

Ethridge, L. E., White, S. P., Mosconi, M. W., Wang, J., Pedapati, E. V., Erickson, C. A., … & Sweeney, J. A. (2017). Neural synchronization deficits linked to cortical hyper-excitability and auditory hypersensitivity in fragile X syndrome. Molecular Autism, 8(1), 22.

Gabard-Durnam, L. J., Mendez Leal, A. S., Wilkinson, C. L., & Levin, A. R. (2018). The Harvard Automated Processing Pipeline for Electroencephalography (HAPPE): standardized processing software for developmental and high-artifact data. Frontiers in neuroscience, 12, 97.

Guggenmos, M., Sterzer, P., & Cichy, R. M. (2018). Multivariate pattern analysis for MEG: A comparison of dissimilarity measures. Neuroimage, 173, 434–447.

Hancock, G. R. (1997). Correlationvalidity coefficients disattenuated for score reliability: A structural equation modeling approach. Educational and Psychological Measurement, 57(4), 598–606.

Hensch, T. K. (2005). Critical period plasticity in local cortical circuits. Nature reviews neuroscience, 6(11), 877–888.

Huttenlocher, P. R. (1979). Synaptic density in human frontal cortex-developmental changes and effects of aging. Brain Res, 163(2), 195–205.

John, E. R., Ahn, H., Prichep, L., Trepetin, M., Brown, D., & Kaye, H. (1980). Developmental equations for the electroencephalogram. Science, 210(4475), 1255–1258.

Klimesch, W. (1997). EEG-alpha rhythms and memory processes. International Journal of psychophysiology, 26(1-3), 319–340.

Lau, Z. J., Pham, T., Chen, S. A., & Makowski, D. (2022). Brain entropy, fractal dimensions and predictability: A review of complexity measures for EEG in healthy and neuropsychiatric populations. European Journal of Neuroscience, 56(7), 5047–5069.

Lippé, S., Kovacevic, N., & McIntosh, R. (2009). Differential maturation of brain signal complexity in the human auditory and visual system. Frontiers in human neuroscience, 3, 792.

McIntosh, A. R., Kovacevic, N., Lippe, S., Garrett, D., Grady, C., & Jirsa, V. (2010). The development of a noisy brain. Archives italiennes de biologie, 148(3), 323–337.

McVoy, M., Lytle, S., Fulchiero, E., Aebi, M. E., Adeleye, O., & Sajatovic, M. (2019). A systematic review of quantitative EEG as a possible biomarker in child psychiatric disorders. Psychiatry research, 279, 331–344.

Merkin, A., Sghirripa, S., Graetz, L., Smith, A. E., Hordacre, B., Harris, R., … & Goldsworthy, M. (2023). Do age-related differences in aperiodic neural activity explain differences in resting EEG alpha? Neurobiology of Aging, 121, 78–87.

Miskovic, V., Ma, X., Chou, C. A., Fan, M., Owens, M., Sayama, H., & Gibb, B. E. (2015). Developmental changes in spontaneous electrocortical activity and network organization from early to late childhood. Neuroimage, 118, 237–247.

Mullen, T. (2012). NITRC: CleanLine: Tool/Resource Inf o. Available online at: http://www.nitrc.org/projects/cleanline (Accessed April 10, 2026).

Paus, T., Zijdenbos, A., Worsley, K., Collins, D. L., Blumenthal, J., Giedd, J. N., … & Evans, A. C. (1999). Structural maturation of neural pathways in children and adolescents: in vivo study. Science, 283(5409), 1908–1911.

Proteau-Lemieux, M., Knoth, I. S., Agbogba, K., Côté, V., Barlahan Biag, H. M., Thurman, A. J., … & Lippé, S. (2021). EEG signal complexity is reduced during resting-state in fragile X syndrome. Frontiers in psychiatry, 12, 716707.

Rhodes, N., Rier, L., Singh, K. D., Sato, J., Vandewouw, M. M., Holmes, N., … & Brookes, M. J. (2025). Measuring the neurodevelopmental trajectory of excitatory-inhibitory balance via visual gamma oscillations. Imaging Neuroscience, 3, imag_a_00527.

Rosseel, Y. (2012). lavaan: An R package for structural equation modeling. Journal of statistical software, 48, 1–36.

Saby, J. N., & Marshall, P. J. (2012). The utility of EEG band power analysis in the study of infancy and early childhood. Developmental neuropsychology, 37(3), 253–273.

Sakai, J. (2020). How synaptic pruning shapes neural wiring during development and, possibly, in disease. Proceedings of the National Academy of Sciences, 117(28), 16096–16099.

Shi, C. T. (2018). Signal pattern recognition based on fractal features and machine learning. Applied sciences, 8(8), 1327.

Uhlhaas, P. J., Roux, F., Rodriguez, E., Rotarska-Jagiela, A., & Singer, W. (2010). Neural synchrony and the development of cortical networks. Trends in cognitive sciences, 14(2), 72–80.

Vakorin, V. A., Lippé, S., & McIntosh, A. R. (2011). Variability of brain signals processed locally transforms into higher connectivity with brain development. Journal of neuroscience, 31(17), 6405–6413.

Walther, A., Nili, H., Ejaz, N., Alink, A., Kriegeskorte, N., & Diedrichsen, J. (2016). Reliability of dissimilarity measures for multi-voxel pattern analysis. Neuroimage, 137, 188–200.

Wang, L. L. (2010). Disattenuation of correlations due to fallible measurement. Newborn and Infant Nursing Reviews, 10(1), 60–65.

Yakovlev, P.I., & Lecours, A. (1967) The myelogenetic cycles of regional maturation of the brain. In: Minkowski, A. (Eds.) Regional Development of the Brain in Early Life (pp. 3–70). Blackwell.

